# Cortical neuron response selectivity derives from strength in numbers of synapses

**DOI:** 10.1101/2019.12.24.887422

**Authors:** Benjamin Scholl, Connon I. Thomas, Melissa A. Ryan, Naomi Kamasawa, David Fitzpatrick

## Abstract

Single neocortical neurons are driven by populations of excitatory inputs, forming the basis of neural selectivity to features of sensory input. Excitatory connections are thought to mature during development through activity-dependent Hebbian plasticity^1^, whereby similarity between presynaptic and postsynaptic activity selectively strengthens some synapses and weakens others^2^. Evidence in support of this process ranges from measurements of synaptic ultrastructure to slice and *in vivo* physiology and imaging studies^3,4,5,6,7,8^. These corroborating lines of evidence lead to the prediction that a small number of strong synaptic inputs drive neural selectivity, while weak synaptic inputs are less correlated with the functional properties of somatic output and act to modulate activity overall^6,7^. Supporting evidence from cortical circuits, however, has been limited to measurements of neighboring, connected cell pairs, raising the question of whether this prediction holds for the full profile of synapses converging onto cortical neurons. Here we measure the strengths of functionally characterized excitatory inputs contacting single pyramidal neurons in ferret primary visual cortex (V1) by combining *in vivo* two-photon synaptic imaging and *post hoc* electron microscopy (EM). Using EM reconstruction of individual synapses as a metric of strength, we find no evidence that strong synapses play a predominant role in the selectivity of cortical neuron responses to visual stimuli. Instead, selectivity appears to arise from the total number of synapses activated by different stimuli. Moreover, spatial clustering of co-active inputs, thought to amplify synaptic drive, appears reserved for weaker synapses, further enhancing the contribution of the large number of weak synapses to somatic response. Our results challenge the role of Hebbian mechanisms in shaping neuronal selectivity in cortical circuits, and suggest that selectivity reflects the co-activation of large populations of presynaptic neurons with similar properties and a mixture of strengths.

## Main

Here we measured visually-driven activity and ultrastructural anatomy of the same synapses on single cortical neurons (Extended Data Figure 1; see Methods). We first performed *in vivo* two-photon imaging of single layer 2/3 pyramidal neurons and dendritic spines on proximal basal dendrites expressing a genetically-encoded activity reporter (GCaMP6s)^9^ to measure functional activity. Following *in vivo* imaging, we perfused the tissue with fixative (see Methods), sectioned tangential to the imaging plane, identified an imaged cell under a confocal microscope, and prepared tissue for serial block-face scanning electron microscopy (SBF-SEM)^10^. After identifying the location of imaged cells within the block (correlating blood vessels and cell bodies), we performed high-resolution SBF-SEM (5.7 - 7 pixels/nm, 56 - 84 nm cutting thickness). We then manually reconstructed the volume of an imaged cell’s soma, dendrites, and visible spines (Fig. 1a). For each spine we reconstructed the spine head, neck, postsynaptic density (PSD), and presynaptic bouton (Fig. 1b). Compared with two-photon images, synaptic ultrastructural anatomy was complex and diverse^11,12^ (Fig. 1b). Visually-driven activity (ΔF/F_o_) from these spines exhibited co-tuning or differential-tuning with respect to the soma (Fig. 1c-d, *right*), as reported previously in ferret V1^9,13^.

**Figure 1:**
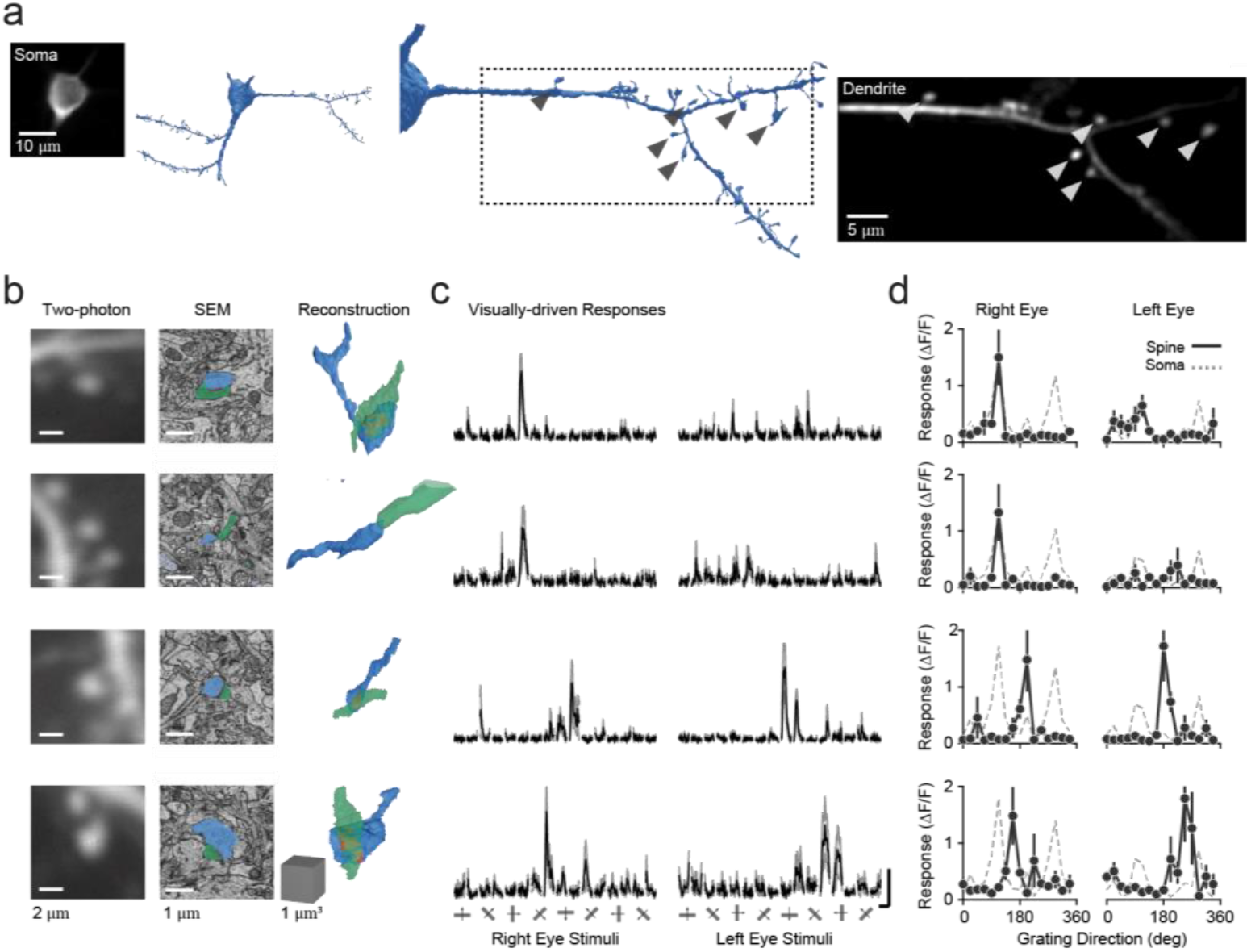
Correlating *in vivo* two-photon synaptic imaging and serial block face scanning electron microscopy. **a**, Example neuron imaged and reconstructed. Shown is a two-photon average-projection of the soma and dendrite. In blue are reconstructions of the soma and dendrites from this cell. Note, only the dendrite and spines are shown. Arrows denote visually-responding spines identified in the EM reconstruction. **b**, Example spines imaged and corresponding EM reconstructions. Shown are a two-photon average-projection *(left column)*, single electron micrograph with annotated synapse components *(middle column),* and the full EM model *(right column)*. **c**, Calcium responses driven by visual stimuli for each spine in (c). Data are mean (black) and standard error (gray). Scale is 100% ΔF/F and 3 sec. **d**, Peak ΔF/F responses across visual stimuli for each spine in (b). Spine data are mean and standard error. Shown also are mean responses of the soma of this cell (gray, dashed line).

We reconstructed 155 visually-responsive (see Methods) synapses imaged *in vivo* on 23 dendritic segments from 5 cells. Most spines (98.7%, n = 153/155) received input from a single presynaptic bouton such that synapses were ‘one-to-one’ connections. A majority (70.0%, n = 109/155) had perforated PSDs, evident by discontinuities in volumetric reconstructions from serial EM sections^14^. Synapse anatomical features varied in size (Spine head volume: mean = 0.39 ± 0.30 μm^3^ s.d., range = 1.26 μm^3^; PSD area: mean = 0.29 ± 0.22 μm^2^ s.d., range = 1.30 μm^2^; Bouton volume: mean = 0.33 ± 0.26 μm^3^ s.d., range = 1.58 μm^3^; Neck length: mean = 1.87 ± 0.86 μm s.d., range = 4.65 μm). Spine head volume and PSD area were strongly correlated, unlike spine head volume and neck length (Extended Data Figure 2)^11^. For comparison with EM features, we calculated functional metrics from peak ΔF/F_o_ responses for each spine (See Methods). Spines exhibited diverse preferences for direction, orientation, and ocular dominance, rather than strictly matching the soma (absolute preference difference ranges = 179.2°, 89.0°, and 1.48 respectively; Extended Data Figure 2). Similarly, there was a wide range in spine selectivity (see Methods) for direction and orientation (0.79 and 0.80, respectively), despite little variation in somatic tuning selectivity (Direction: 0.21 ± 0.09, Orientation: 0.48 ± 0.04, Mean ± SE). In sum, the populations of synaptic inputs on individual cells exhibit both a wide range of strengths (small and large) and functional properties (aligned and misaligned to the soma), raising the question whether there is a systematic functional synaptic weight distribution relating structure and function.

To test whether synaptic strength is functionally-biased, whereby strong synaptic inputs drive neural selectivity and are co-tuned to the somatic output, we directly compared synapse structural and functional properties. For simplicity, we first focus on orientation preference. Surprisingly, the strength of individual synapses was uncorrelated with functional similarity to the somatic output (i.e. absolute orientation preference difference) (Fig. 2). We found no relationship for spine head volume (Circular-linear r = 0.03, p = 0.91) or PSD area (Fig. 2a-b; Circular-linear r = 0.12, p = 0.34). In particular, and in contrast to a recent study^7^, PSD size was not significantly different between co-tuned (Δθ < 45°, n = 100) and orthogonally-tuned (Δθ > 45°, n = 57) spines (p = 0.72). Similarity in orientation preference and PSD area did not depend on spine distance from the soma (< 50 μm: p = 0.99, n = 77; > 50 μm: p = 0.84, n = 78). As PSD area is a structural correlate of synaptic strength^15,16^ and spine necks can attenuate presynaptic drive^17^, we generated a separate metric comprised of all anatomical features using a NEURON model^18^ (see Methods). For each synapse we simulated depolarization in spine and soma (compartments after a single presynaptic spike (Extended Data Figure 2). Even for simulated somatic depolarization, we uncovered no systematic relationship (Fig. 2c; Circular-linear r = 0.08, p = 0.60). Null relationships were also found for direction preference, ocular dominance, and spine-soma tuning correlation (Extended Data Fig. 3). To ensure our results are accurate, we tightened inclusion criteria and performed the same analyses. For spines exhibiting low residual-correlation with global dendritic events (r < 0.2, n = 75) or high signal-to-noise ratio (SNR > 3, n = 71) we uncovered no correlation between any functional or anatomical property examined (Extended Data Figs. 4-5). Even when analyzing spine populations from individual cells, we consistently observed no correlations (Extended Data Fig. 6). These data suggest that strong and weak synapses on individual neurons are equally likely to be functionally similar or dissimilar to the somatic output.

**Figure 2:**
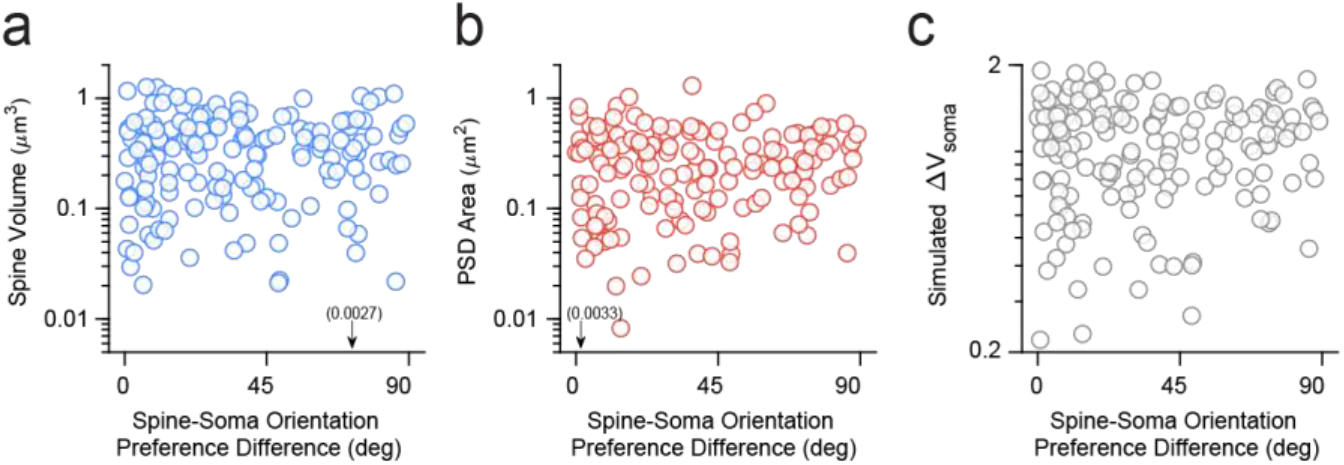
Similarity in spine-soma orientation preference is uncorrelated with synaptic strength. **a**, Difference in orientation preference between spine and corresponding soma and the spine head volume of that spine. Each data point *(blue)* represents an individual visually-responsive synapse reconstructed (n = 155 from 5 cells). Gray data points are binned averages. Note the ordinate on a log scale. Arrow denotes a single data point outside the ordinate limits. **b**, Same as in (a) for PSD area *(red).* **c**, Same as in (a) for NEURON simulation of somatic depolarization for each spine reconstructed *(gray)* correlated. Note ordinate scale is linear.

How might synaptic populations contribute to neural selectivity without functionally-biased synaptic weight distributions? We propose that a major factor contributing to selectivity is the total number of synapses recruited. To examine this possibility we compared synaptic aggregate predictions with the average somatic orientation tuning across our 5 cells, focusing on dominant-eye stimuli. As spine ΔF/F_o_ does not reflect strength^19^ and PSD area was uncorrelated with maximum response amplitude (Spearman’s r = 0.07, p = 0.19), we converted spine ΔF/F_o_ into discrete calcium events (see Methods). We defined ‘active’ synapses as those with calcium events in at least 50% of trials of a particular stimulus and calculated the total weight by summing PSD area across active synapses for each stimulus orientation (±67.5° around somatic preference). Total average synaptic weight was selective for the somatic preferred orientation (Fig. 3a), however, a main determinant for this selectivity was the *total number* of active synapses contributing to each stimulus condition (Fig. 3a-b), which were normally distributed about the somatic preference (p = 0.36, Lilliefors test). No differential recruitment of strength was evident across stimuli (Fig. 3b; p = 0.52, Kruskal-Wallis test). These observations were consistent for the majority of cells and respective synaptic populations in this study (n = 4/5): Active synapses were preferentially recruited around the somatic-preferred stimulus (p ≥ 0.37, Lilliefors test) and we found no difference in strength across stimuli (p ≥ 0.31, Kruskal-Wallis test). Thus, synaptic aggregate tuning is due, in part, to an overrepresentation of somatic preference in synaptic populations (Extended Data Figure 2), which leads to increased numbers of synapses recruited for preferred stimuli. Not surprisingly, the overrepresentation of somatic preference in input populations, diminishes the impact that functional biases in the strength of individual synapses can have on aggregate input selectivity (Extended Data Figure 7). Thus, our data do not support the hypothesis that the strongest synaptic inputs contribute disproportionally to somatic selectivity. Instead, our data reveal a potentially more important factor: the total number of individual synapses recruited by sensory stimuli.

**Figure 3:**
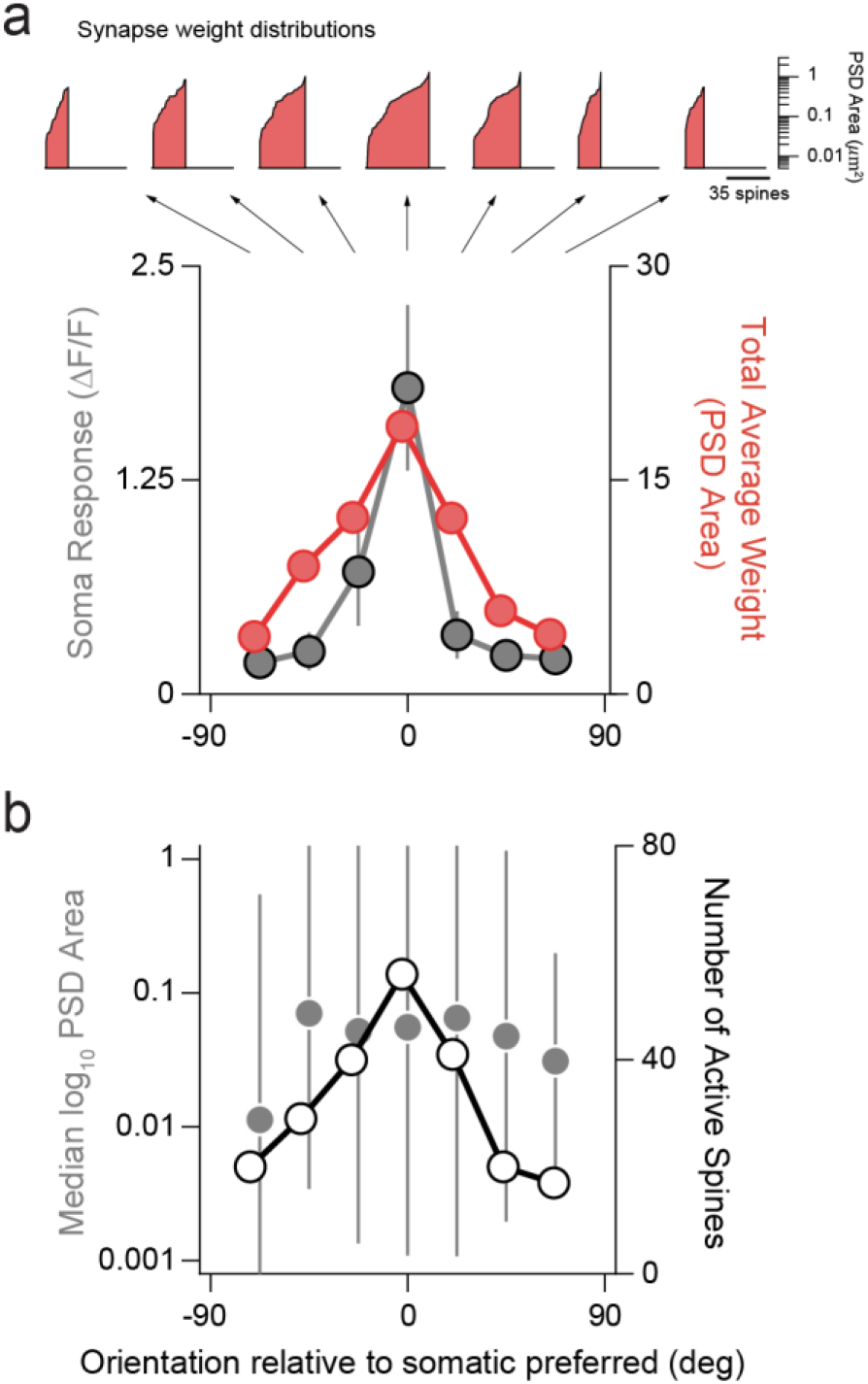
Somatic selectivity predicted by total weight derived from total number active synapses. **a**, Average somatic orientation tuning (n = 5 cells, *gray*) compared to the total average weight (summed PSD area, *red)* from active synapses for each stimulus condition (θ_soma pref_ ± 67.5°). Active synapses defined as those exhibiting calcium events on at least 50% of stimulus presentation trials. Soma data are mean and standard error. Total average weight data are summed PSD area across active spines. Shown at top are cumulative distributions of PSD area for active synapses for each stimulus condition. Note, ordinate is PSD area (μm^2^) and abscissa is total number of active spines. **b**, Plots of median log_10_ PSD area and interquartile range across active spines (gray circles) and total number of active spines (white circles) for each orientation. Note, data are derived from distributions shown in **a**.

Another factor contributing to neural selectivity is spatial clustering of co-active synaptic inputs. Combined strength of co-active synapses is enhanced by spatial proximity^20^, and functionally-similar, neighboring co-active synapses^9^ are proposed to exhibit greater strength through cooperative plasticity^21^. It is unknown, however, whether synaptic clustering relates to strength. We first computed distance-dependent trial-to-trial correlation between pairs of synapses, replicating a trend reported previously^9^ (Fig. 4a). We then restricted pairwise comparisons to either weak (spine volume < 0.35 μm^3^) or strong (spine volume > 0.35 μm^3^) synapses. Functional clustering at small distances was only evident for weaker synapses, not strong synapses (Fig. 4b-c). In addition, functional clustering of weaker inputs persisted when examining only spines with a similar orientation preference as their corresponding soma (Fig. 4d). Repeating the same analyses after removing stimulus trials containing a dendritic calcium event (see Methods), produced similar results (Extended Data Figure 8). We also extended these analyses to synapse pairs with short (<1.75 μm) or long (>1.75 μm) spine necks, but found no differences in distance-dependent correlations between groups (Extended Data Figure 8). Taken together, these analyses suggest that larger (stronger) synapses are more spatially-isolated in activity and the spatiotemporal clustering of smaller (weaker) synapses might act to enhance their combined synaptic strength in numbers.

**Figure 4:**
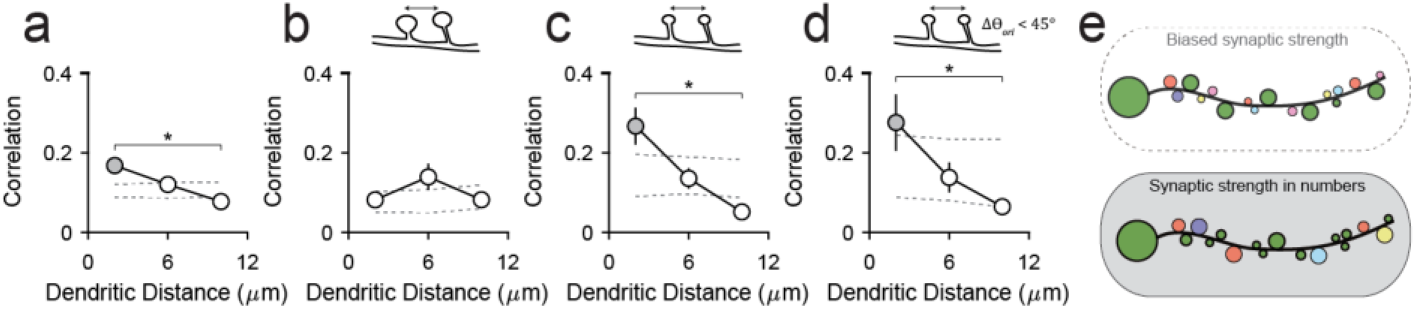
Smaller, but not larger, synapses exhibit local spatiotemporal clustering. **a**, Relationship between spine pair distance and trial-to-trial correlation during visual stimulation. Data are mean and SEM *(black).* Also shown is shuffled correlations *(gray dashed lines),* data are SEM. Gray data points denote those significantly greater than the shuffled correlations (p < 0.05, bootstrapped confidence interval). Asterisk denotes significantly different correlation distributions (p < 0.05, Mann-Whitney test). **b**, Same as in (a) for large (spine volume > 0.35 μm^3^) synapse pairs. **c**, Same as in (a) for small (spine volume < 0.35 μm^3^) synapse pairs. **d**, Same as in (a) for small (spine volume < 0.35 μm^3^) synapse pairs with orientation preference similar to the somatic output (Δθ < 45°). **e**, Illustration of two competing models of functional synaptic strength. Our data do not support functionally-biased synaptic strength *(top)*. Instead, our data suggest strength in numbers *(bottom)* whereby inputs co-tuned with the somatic are more numerous but exhibit a wide range in strengths.

Correlating *in vivo* synaptic imaging and EM, to measure functional properties and anatomical strength, we tested a hypothesis that strong synaptic inputs drive neural selectivity, while weak synaptic inputs are not structured and act to modulate activity overall^6,7^. We found no evidence to support this hypothesis. Instead our data suggest, simply, that activation of a greater number of synapses equates to greater synaptic weight overall (Fig. 4e). Weaker synapses are greater in number overall as synaptic strength is lognormally distributed^22^ and spatial clustering may act to enhance their effect on somatic activity. Notably, we likely underestimate the total number of weak synapses contributing to the somatic output. *In vivo* two-photon microscopy captures larger dendritic spines and back-propagating action potentials likely mask weaker spines co-active with the soma.

One possible explanation of our observations is that postsynaptic spiking activity shapes the overall distribution of functional properties of input populations during plasticity, rather than modulating individual synapse strength. This would give rise to the soma-biased input populations, as observed in this study and others^9,13^. This could be achieved by modulating synaptogenesis and synaptic pruning^23^; increasing the probability of stabilizing inputs co-tuned with the somatic output. While developmental models of single neurons have made similar predictions^24^, this process has yet to be observed *in vivo.* Unitary synaptic strength, instead, may simply depend on operational properties of presynaptic afferents. In fact, when comparing spine selectivity to motion direction or orientation with anatomical correlates of strength we did find significant correlations (Extended Data Figure 9). Thus, synaptic strength may reflect the reliability of afferents representing particular visual features or the afferent dynamic range in spike rate. Synaptic strength in this case might not follow Hebbian spike-timing-dependent-plasticity^2^, but instead, depend on non-Hebbian plasticity^25^ or local signaling mechanisms^26^.

Why do our measurements differ from previous studies? First, no previous study has assessed the functional properties and strength of a population of excitatory synapses that converge onto a single cortical neuron. Synaptic strength has conventionally been defined by somatic EPSP amplitude recorded with electrophysiology^27–29^. Somatic EPSPs likely result from both the number^27–29^, strength of presynaptic contacts^30^, and the distance of those contacts from the soma. Further, the difficulty of such measurements leads to a bias for identifying connections between nearby neurons, as opposed to our more unbiased approach of imaging dendritic spines, whose presynaptic partner may reside locally or project long-range. Local sampling could explain the different conclusion derived from experiments that used correlative EM and cellular imaging to assess synaptic inputs from nearby layer 2/3 neurons^7^. Also, most previous studies probed circuit organization in the mouse and we cannot exclude the possibility that there are fundamental differences in circuit design between rodents and carnivores. Clearly, there is more to be learned about the synaptic weight distributions of cortical neurons, including whether they vary for different sources of inputs, and different dendritic compartments^31^. These results make it clear that the significance of synaptic weight lies in factors beyond response selectivity, and challenge prevailing views of the developmental mechanisms that shape this selectivity.

