## Supplementary material for "Cortical neuron response selectivity derives from strength in numbers of synapses": Methods

All procedures were performed according to NIH guidelines and approved by the Institutional Animal Care and Use Committee at Max Planck Florida Institute for Neuroscience.

*Viral Injections*

Female ferrets (n = 3) aged P18-23 (Marshall Farms) were anesthetized with isoflurane (delivered in O_2_). Atropine was administered and a 1:1 mixture of lidocaine and bupivacaine was administered SQ. Animals were maintained at an internal temperature of 37^o^ Celsius. Under sterile surgical conditions, a small craniotomy (0.8 mm diameter) was made over the visual cortex (7-8 mm lateral and 2-3 mm anterior to lambda). A mixture of diluted AAV1.hSyn.Cre (1:25000 to 1:50000) and AAV1.Syn.FLEX.GCaMP6s (UPenn) was injected (125 - 202.5 nL) through beveled glass micropipettes (10-15 µm outer diameter) at 600, 400, and 200 µm below the pia. Finally, the craniotomy was filled with sterile agarose (Type IIIa, Sigma-Aldrich) and the incision site was sutured.

*Cranial Window*

After 3-5 weeks of expression, ferrets were anesthetized with 50mg/kg ketamine and isoflurane. Atropine and bupivacaine were administered, animals were placed on a feedback-controlled heating pad to maintain an internal temperature of 37° Celsius, and intubated to be artificially respirated. Isoflurane was delivered throughout the surgical procedure to maintain a surgical plane of anesthesia. An intravenous cannula was placed to deliver fluids. Tidal CO_2_, external temperature, and internal temperature were continuously monitored. The scalp was retracted and a custom titanium headplate adhered to the skull (Metabond, Parkell). A craniotomy was performed and the dura retracted to reveal the cortex. One piece of custom coverglass (3 mm diameter, 0.7 mm thickness, Warner Instruments) adhered to a custom insert using optical adhesive (71, Norland Products) was placed onto the brain to dampen biological motion during imaging. A 1:1 mixture of tropicamide ophthalmic solution (Akorn) and phenylephrine hydrochloride ophthalmic solution (Akorn) was applied to both eyes to dilate the pupils and retract the nictitating membranes. Contact lenses were inserted to protect the eyes. Upon completion of the surgical procedure, isoflurane was gradually reduced and pancuronium (2 mg/kg/hr) was delivered IV.

*Visual Stimuli*

Visual stimuli were generated using Psychopy^32^. The monitor was placed 25 cm from the animal. Receptive field locations for each cell were hand mapped and the spatial frequency optimized (range: 0.04 to 0.20 cpd). For each soma and dendritic segment, square-wave or sine-wave drifting gratings were presented at 22.5 degree increments to each eye independently (2 second duration, 1 second ISI, 8-10 trials for each field of view). Drifting gratings of different directions (0 – 315^o^) were presented independently to both eyes.

*Two-photon imaging*

Two-photon imaging was performed on a Bergamo II microscope (Thorlabs) running Scanimage^33^ (Vidrio Technologies) with 940 nm dispersion-compensated excitation provided by an Insight DS+ (Spectraphysics). For spine and axon imaging, power after the objective was limited to <50 mW. Cells were selected for imaging on the basis of their position relative to large blood vessels, responsiveness to visual stimulation, and lack of prolonged calcium transients resulting from over-expression of GCaMP6s. Images were collected at 30 Hz using bidirectional scanning with 512x512 pixel resolution or with custom ROIs (region of interest; framerate range: 22 - 50 Hz). Somatic imaging was performed with a resolution of 2.05 - 10.24 pixels/ µm. Dendritic spine imaging was performed with a resolution of 6.10 -15.36 pixels/µm.

*Two-Photon Imaging Analysis*

Imaging data were excluded from analysis if motion along the z-axis was detected. Dendrite images were corrected for in-plane motion via a 2D cross-correlation based approach in MATLAB or using a piecewise non-rigid motion correction algorithm^34^. ROIs were drawn in ImageJ; dendritic ROIs spanned contiguous dendritic segments and spine ROIs were fit with custom software. Mean pixel values for ROIs were computed over the imaging time series and imported into MATLAB^35^. $\Delta F/{F_{o}}$was computed by computing $F_{o}$ with time-averaged median or percentile filter (10th percentile). For spine signals, we subtracted a scaled version of the dendritic signal to remove back-propagating action potentials as performed previously^13^. $\Delta F/{F_{o}}$ traces were synchronized to stimulus triggers sent from Psychopy and collected by Spike2.

Peak $\Delta F/{F_{o}}$ responses to bars and gratings were computed using the Fourier analysis to calculate mean and modulation amplitudes for each stimulus presentation, which were summed together. Spines were included for analysis if the mean peak $\Delta F/{F_{o}}$for the preferred stimulus was >10% $\Delta F/{F_{o}}$, the SNR^9^ at the preferred stimulus was > 1, and spines were weakly correlated with the dendritic signal (Spearman’s correlation, r < 0.4). Some spine traces contained negative events after subtraction, so correlations were computed ignoring negative $\Delta F/{F_{o}}$ values. Preferred orientation and direction for each spine was calculated by fitting responses with a double Gaussian tuning curve^13^ using lsqcurvefit (Matlab). Ocular dominance index was calculated as the normalized difference between preferred left and right eye responses^36^. Spine-soma tuning correlation was computed as the Pearson's correlation (Matlab) between mean responses a spine and mean responses of the somatic output. Orientation and direction selectivity was computed by calculating the vector strength of mean responses^37^. For local clustering analyses, trial-to-trial correlations were computed as the correlation of peak ΔF/F responses to each stimulus on a per-trial basis. To identify spine or dendritic calcium events, ΔF/F traces were smoothed with an exponentially weighted moving average filter (MATLAB) and locating the peaks of calcium events. Peak amplitude of calcium events were compared to the standard deviation of baseline spine fluorescence values prior to subtraction. Spines used for analysis of spontaneous activity were required to have at least 1 calcium event (peak amplitude > 3 s.d. background fluorescence) independent of dendritic calcium events.

*NEURON Modeling*

For each synapse reconstructed, we simulated the change in membrane potential at the spine and soma due to a single action potential arriving at the synapse (on the spine head). Simulations of anatomical features allow generation of a single metric (voltage attenuation between spine head and soma) accounting for a variety of synapse features. We modeled a somatic compartment (radius = 13 μm, *g_Na_* = 0 S/cm^2^, *g_k_* = 0.036 S/cm^2^, *g_leak_* = 0.003 S/cm^2^, *E_leak_* = -50 mV, *R_a_* = 105 Ωcm, *C_m_* = 1 μF/cm^2^) connected to a 400 μm long dendrite (diameter = 1 μm, *R_a_* = 105 Ωcm, *C_m_* = 1 μF/cm^2^). Each spine was placed on the dendrite at the distance from soma as measured with EM and connected via a ‘neck’ to a ‘spine head’ where a synapse was placed. Synapse compartments had the same basic properties (*R_a_* = 250 Ωcm, *C_m_* = 1 μF/cm^2^) and passive conductances. Spine neck diameter was fixed to 200 nm, matching widths measured in serial EM sections (data not shown) and the length was set to measured values. Spine head length was set to 1 μm so the diameter could be determined from volume measurements (assigning spine heads to be a cylindrical compartment):

$$D=\sqrt{V/{4\pi}}$$

Next, we converted measured PSD area into a value describing the max synaptic conductance. Here we make several assumptions. Based on the linear correlation between the number of receptors and PSD size, we approximate ~0.87 receptors and ~2.0 receptors per 100 nm for AMPA and NMDA, respectively^38^. As a simplification, we extract PSD diameter as if our PSDs were circular (as above). Then an AMPA conductance is

$$g_{AMPA}=D_{spine}\cdot\left( {0.87}/{0.100} \right)\cdot g_{R}$$

where *g_R_* is 15 pS per channel. In this way, measured PSD area is linearly related to the synaptic conductance used in each model. For each simulation, parameters were set and the maximum depolarization from *V_rest_* (-67.5 mV) was measured in the somatic and spine head compartment.

In this paper we only show simulations of AMPA conductance, but we also ran simulations with an additional NMDA conductance.

Simulated voltage attenuation (ΔVm_soma_/ ΔVm_spine_) for synapses were strongly correlated (r = 0.99, p < 0.001, Spearman’s correlation) so we expect our results to be the same for either model.

*Serial Block-Face Scanning Electron Microscopy (SBF-SEM)*

Five layer 2/3 pyramidal neurons from 3 animals previously imaged with *in vivo* two-photon microscopy were imaged with SBF-SEM. A total of 23 segments of proximal basal dendrites and 155 spines were reconstructed and analyzed. To facilitate EM reconstruction we limited imaging to proximal basal dendrites.

Fixed (2% paraformaldehyde and 2% glutaraldehyde in a 0.1 mM sodium cacodylate) brain slices of 80 µm thickness were trimmed to approximately 2 × 2 mm to contain the cell of interest at the center. This was accomplished by using blood vessels and slice edges, visible in a 20x epifluorescence image of the slice, as landmarks. The tissue pieces were incubated in an aqueous solution of 2% osmium tetroxide buffered in 0.1 mM sodium cacodylate for 45 minutes at room temperature (RT). Tissue was not rinsed and the osmium solution was replaced with cacodylate buffered 2.5% potassium ferrocyanide for 45 minutes at RT in the dark. Tissue was rinsed with water 2 x 10 minutes, which was repeated between consecutive steps. Tissue was incubated in warm (60˚C) aqueous 1% thiocarbohydrizide for 20 minutes, aqueous 1% osmium tetroxide for 45 minutes, and then 1% uranyl acetate in 25% ethanol for 20 minutes. Tissue was rinsed then left in water overnight at 4˚C. The following day, tissue was stained with Walton’s lead aspartate for 30 minutes at 60˚C. Tissue was then dehydrated in a graded ethanol series (30, 50, 70, 90, 100%), 1:1 ethanol to acetone, then 100% acetone. Tissue was infiltrated using 3:1 acetone to Durcupan resin (Sigma Aldrich) for 2 hours, 1:1 acetone to resin for 2 hours, and 1:3 acetone to resin overnight, then flat embedded in 100% resin on a glass slide and covered with an Aclar sheet at 60˚C for 2 days. Since SBF-SEM requires conductive samples to minimize charging during imaging, the tissue was trimmed to less than 1 × 1 mm and one side was exposed using an ultramicrotome (UC7, Leica), then turned downwards to be remounted to a metal pin with conductive silver epoxy (CircuitWorks, CHEMTRONICS).

Tissue was sectioned and imaged using 3View and Digital Micrograph (Gatan Microscopy Suite) installed on a Gemini SEM300 (Carl Zeiss Microscopy LLC.) equipped with an OnPoint BSE detector (Gatan, Inc.). The detector magnification was calibrated within SmartSEM imaging software (Carl Zeiss Microscopy LLC.) and Digital Micrograph with a 500 nm cross line grating standard. A low magnification image of each block face was matched to its corresponding depth in the confocal Z-stack in Adobe Photoshop (CS6 version 13.0.1) using blood vessels and cell bodies as fiducials. These features were clear across magnification scales (from 10x to ~10,000x) and used to estimate the XY position and depth of the cell and proximal segments of basal dendrites. Final imaging was performed at 2.0-2.2 kV accelerating voltage, 20 or 30 µm aperture, working distance of ~5 mm, 0.5-1.2 µs pixel dwell time, 5.7-7 nm per pixel, knife speed of 0.1 mm/sec with oscillation, and 56 - 84 nm section thickness. Imaged volumes ranged from 125x125x36 µm to 280x170x52 µm. Serial images were exported as TIFFs to TrakEM2^39^ and aligned using Scale-Invariant Feature Transform image alignment with linear feature correspondences and rigid transformation^40^. Once aligned, each dendrite of interest was cropped from the full volume to reduce computational overhead in subsequent analyses. Aligned images were exported to Microscopy Image Browser^41^ for segmentation of dendrites, spines, PSDs, and boutons. Following thin spine necks at our Z resolution was sometimes impossible, or in some cases a single spine imaged with the light microscope was actually resolved as several in EM.

Binary labels files were then imported to Amira (versions 6.7, 2019.1) which was used to create 3D surface models of each dendrite, spine, PSD, and bouton. Once reconstructed, each reconstructed dendrite was overlaid onto its corresponding two-photon image using Adobe Photoshop for re-identification of individual spines. Amira was used to measure the volume of spine heads and boutons, surface area of PSDs, and spine neck length. Finally, Blender (versions 2.79, 2.8) was used to create 3D renderings.

In these data, presynaptic boutons connected at most 2 different spine heads, and was observed for 12% of synapses examined (n = 25/202). Of all spines imaged *in vivo*, 57% (n = 167/292) could be re-identified in EM volumes. Reconstructed spines resolved *in vivo* were typically larger than those unresolvable under two-photon excitation or unresponsive (mean = 0.46 ± 0.29 μm^3^ s.d., n = 38, 0.28 ± 0.29 μm^3^ s.d., n = 20, respectively; p = 0.014).

*Statistics*

Statistical analyses are described in the main text and in figure legends. We used non-parametric statistical analyses (Wilcoxon sign-rank test, rank-sum test, Kruskal-Wallis test) or permutation tests to avoid assumptions about the distributions of the data. All statistical analysis was performed in MATLAB. Circular-linear correlation coefficients were computed for tests with circular variables (orientation and direction preference). For all other tests, Spearman’s correlation coefficient was computed. All correlation significance tests were one-sided. Lilliefors test for normality was used on the data presented in Figure 3. Quantitative approaches were not used to determine if the data met the assumptions of the parametric tests.

**Data and code availability**

Data and code are available from the corresponding author upon reasonable request.

**Acknowledgements**

The authors thank Clara Tehpol for surgical assistance, David Hildebrand for discussion of SEM techniques, Balazs Ujfalussy for help with NEURON modeling, Nicole Shultz and Rachel Satterfield for help with perfusions and fixative preparation, the Fitzpatrick lab for useful discussions, and the MPFI ARC for animal care. The authors thank the GENIE project for access to GCaMP6.

**Author contributions**

B.S. conceived experiments. B.S. performed biological experiments. C.T. and M.R. preformed electron microscopy and image processing with guidance from N.K.. C.T., M.R., and B.S. preformed volumetric reconstruction. B.S. analyzed data with guidance from D.F.. B.S. wrote the paper with help from C.T., M.R., and D.F..

**Competing financial interests**

The authors declare no competing financial interests.

**Methods References**

32.	Peirce, J. W. PsychoPy--Psychophysics software in Python. *J. Neurosci. Methods* **162,** 8–13 (2007).

33.	Pologruto, T. A., Sabatini, B. L. & Svoboda, K. ScanImage: flexible software for operating laser scanning microscopes. *Biomed Eng Online* **2,** 13 (2003).

34.	Pnevmatikakis, E. A. & Giovannucci, A. NoRMCorre: An online algorithm for piecewise rigid motion correction of calcium imaging data. *J. Neurosci. Methods* **291,** 83–94 (2017).

35.	Sage, D., Prodanov, D. & Tinevez, J. Y. MIJ: making interoperability between ImageJ and Matlab possible. *bigwww.epfl.ch* (2012).

36.	Scholl, B., Pattadkal, J. J., Dilly, G. A., Priebe, N. J. & Zemelman, B. V. Local integration accounts for weak selectivity of mouse neocortical parvalbumin interneurons. *Neuron* **87,** 424–436 (2015).

37.	Scholl, B., Tan, A. Y. Y., Corey, J. & Priebe, N. J. Emergence of orientation selectivity in the Mammalian visual pathway. *J. Neurosci.* **33,** 10616–10624 (2013).

38.	Takumi, Y., Ramírez-León, V., Laake, P., Rinvik, E. & Ottersen, O. P. Different modes of expression of AMPA and NMDA receptors in hippocampal synapses. *Nat. Neurosci.* **2,** 618–624 (1999).

39.	Cardona, A. *et al.* TrakEM2 software for neural circuit reconstruction. *PLoS One* **7,** e38011 (2012).

40.	Lowe, G. SIFT-the scale invariant feature transform. *Int. J* (2004).

41.	Belevich, I., Joensuu, M., Kumar, D., Vihinen, H. & Jokitalo, E. Microscopy image browser: A platform for segmentation and analysis of multidimensional datasets. *PLoS Biol.* **14,** e1002340 (2016).
